# Low genetic diversity in a population of Tricolored Blackbird (*Agelaius tricolor*), a species pending Endangered status

**DOI:** 10.1101/2020.07.13.201574

**Authors:** Irene A. Liu, Robert J. Meese

## Abstract

The Tricolored Blackbird (*Agelaius tricolor*) is a colonial songbird, found almost exclusively in California, whose total population size has sharply declined over the past century. It is currently under review to be listed as Endangered under both the California and U.S. Endangered Species Acts. Here we assess the genetic diversity of a breeding population in California’s Central Valley, comparing our findings with previously sampled conspecific and congeneric populations. First, we genotyped 50 adults at 9 microsatellite loci in our focal population and estimated allelic and Shannon diversity, observed and expected heterozygosity, and the inbreeding coefficient (F_IS_). Second, we compared our results to those of the one existing study on Tricolored Blackbird conservation genetics and found that levels of allelic diversity and heterozygosity in our focal population were similar to those of 11 previously studied populations. Unlike the earlier study, which found moderately high mean inbreeding coefficients, we detected no evidence of inbreeding in our focal population. Third, we used 7 of the 9 loci to compare the genetic diversity of our focal population with populations of 2 previously sampled *Agelaius* congeners. We found that allelic diversity, Shannon diversity, and expected heterozygosity in our Tricolored Blackbird population were most similar to those of an isolated Red-winged Blackbird (*A. phoeniceus*) population in the Bahamas. We discuss possible reasons for the different results from the conspecific study, outline why the collective findings from both studies support the need for protective measures, and urge conservation action to maintain existing genetic diversity and gene flow before ongoing population losses lead to adverse fitness consequences.

## INTRODUCTION

The Tricolored Blackbird (*Agelaius tricolor*) is a songbird that nests in colonies restricted to lower-elevation locations of California, with small remnant populations in Oregon, northern Baja California, and western Nevada and Washington (Meese et al. 2014). Once abundant throughout California’s Central Valley, this species has declined sharply in total population size over the last century and especially in recent decades (Beedy 2008; Meese et al. 2014; Meese 2015). Colonies of 100,000 or more Tricolored Blackbirds were common from the 1920s to 2006 (Neff 1937; Orians 1961; Payne 1969; Meese 2006), but in 2013 and 2014 the average size of the largest colony was ∼25,000 birds (Meese 2015). While different census and documentation methods make cross-study estimates of total population size difficult to compare (Meese 2014), within-study surveys report steady and steep declines (e.g., 1930s-1970s, DeHaven et al. 1975; 1992-2002, Cook and Toft 2005). Standard estimation methods implemented in 2008 showed that statewide population estimates of Tricolored Blackbirds fell from 400,000 in 2008 to 145,000 birds in 2014 (Meese 2014).

These declines are attributed to 2 primary causes: the widespread loss of the Tricolored Blackbird’s native wetland habitat (estimated at a 96% loss over the last 150 years, Kreissman 1991) to agriculture and urbanization (Beedy 2008; Meese et al. 2014), and the species’ switch to nesting in active grain fields that are harvested as part of normal agricultural operations, which has resulted over several decades in complete losses of many of the largest colonies. Because of land-use changes and fragmentation of suitable habitat, breeding site occupancy of Tricolored Blackbirds has declined 3 times as quickly as sites have been recruited (Holyoak et al. 2014).

Additional factors include low insect abundance and the predominance of a grain diet in grain-field-nesting colonies, leading to inadequate nutrition and decreased reproductive success (Meese 2013). Low insect abundance may be related to the widespread use of pesticides, especially of neonicotinoids, which are applied to croplands surrounding nesting colonies and can significantly impact insectivorous bird populations (Hallmann et al. 2014). Moreover, predation levels often are high and can be human-mediated, such as when lowering of water levels in managed wetlands enables access by terrestrial predators (Beedy 2008). Post-breeding birds foraging in rice fields also are shot when in mixed-species flocks containing the morphologically similar Red-winged Blackbird (*A. phoeniceus*), which is permitted to be killed as an agricultural pest in California (Meese 2015). Alongside all of these threats, the species’ colonial nature means that it is prone to acute and large-scale losses.

In response to such threats, the Tricolored Blackbird was recognized as Endangered by the IUCN Red List in 2006. In California, the blackbird has been listed as a Species of Special Concern since 1990. The California Department of Fish and Wildlife granted the species emergency protection under the California Endangered Species Act (CESA) in December 2014 but 6 months later declined to renew these protections. Following a petition re-submission (Belenky and Bond 2015), as of December 2015 the species is again under protection and has advanced to candidacy to be listed under the CESA. Federally, the status of the Tricolored Blackbird is currently under review for possible listing under the U.S. Endangered Species Act (ESA).

Despite decades of study on the Tricolored Blackbird’s demography and natural history, limited data exist for the species’ genetic profile. To our knowledge, 2 studies have profiled molecular patterns in Tricolored Blackbirds. The first study was a conservation genetics analysis using data from 8 microsatellites and 2 mitochondrial genes (Berg et al. 2010). The authors found evidence of gene flow within and across 7 populations in the Central Valley, sampled from 2001-2005, and 4 in southern California, sampled from 2007-2008. Both regions additionally had moderately high mean inbreeding coefficients (F_IS_ = 0.121 and 0.090, respectively). Because of the lack of population structure, the authors concluded that Tricolored Blackbirds in different areas did not need to be considered as separate management units. The second study (Barker et al. 2012) used mitochondrial DNA from 10 Tricolored Blackbirds from a southern California population and 31 Red-winged Blackbirds from multiple sources to compare information content in ND2 vs. the control region. Although the study’s intent was to contrast evolutionary patterns in coding vs. non-coding regions, the results nevertheless revealed that Tricolored Blackbirds had lower genetic diversity than Red-winged Blackbirds.

Here we use microsatellite data to estimate levels of genetic diversity and inbreeding in a separate breeding population in the Central Valley. Because of the limited inference that can be drawn from a single-population study, we present our results alongside previously obtained profiles of conspecific and congeneric populations. We compare our estimates of allelic diversity, observed and expected heterozygosity, and the inbreeding coefficient with those in Berg et al. (2010)’s conservation genetics analysis of 11 Tricolored Blackbird populations. We also compare allelic diversity, Shannon diversity, and expected heterozygosity with previously studied populations of 2 *Agelaius* congeners, Red-winged Blackbirds (*A. phoeniceus*) and Yellow-Shouldered Blackbirds, (*A. xanthomus*), that span a gradient of genetic diversity (Liu 2015; Liu et al. 2015).

## METHODS

### Field Sampling

We collected blood samples from 8 males and 42 females from 42 nests in a colony breeding in a manmade cattail pond on the Conaway Ranch, Woodland, CA (coordinates: 38°38’49.2”N, 121°42’07.2”W). Sampling took place from June 5 to 17, 2013. Except for 2 males caught opportunistically outside the pond, adults were captured using walk-in traps placed over their nests. This method was the only way to identify individuals to their territories. Territory could not be assigned by behavior, as adults were closely spaced and flushed collectively from their nests when an observer approached. The low number of captured males is due to their lower provisioning rates and, when they did approach the nest, their greater aversion to entering traps (I.A. Liu personal observation).

Sampling methods followed Liu et al. (2015): Adults were bled from the brachial vein using sterile 26G 9 ½ in. BD PrecisionGlide needles, blood was collected onto Whatman FTA bloodstain cards treated with 1 M EDTA, and adults were banded with USFWS and color bands.

### DNA Extraction and Genotyping

We genotyped all individuals at 9 microsatellite loci (Table S1) following DNA amplification with the PCR profile described in Liu (2015). Of these loci, 5 were derived from Red-winged Blackbird (Barker et al. 2011) and verified not to show ascertainment bias relative to the other 4 loci, originally identified in more distantly related species (Liu 2015). Plates were processed at Eton Bioscience Inc., and genotypes were scored with GeneMarker 1.8 (SoftGenetics) using size standard GS-500.

Homozygous alleles were genotyped at least twice to account for allelic dropout. We further checked for dropout and null alleles using Micro-Checker 2.0 (Van Oosterhout et al. 2004) for all but 4 loci (LTMR6, Ap107, Ap144, and Dpµ16), which had irregular alleles outside the base-pair lengths expected from the motif. We tested all loci for Hardy-Weinberg equilibrium and pairwise linkage disequilibrium using Genepop (Raymond and Rousset 1995).

### Genetic Diversity

We first used GenAlEx 6.501 (Peakall and Smouse 2012) to calculate mean allelic diversity, Shannon diversity, observed and expected heterozygosity, and the inbreeding coefficient (the fixation index F_IS_) for all loci for the 50 adults. Values reported are mean ± SE.

Next, we compared measures of genetic diversity between the Conaway Ranch Tricolored Blackbird population and the 11 Tricolored Blackbird populations in Berg et al. (2010), particularly because only one locus (Dpµ16) was shared between the 2 studies. We performed an ANOVA with raw allelic diversity and the observed and expected heterozygosity per locus across all 3 groups (the Conaway Ranch population, 4 southern California populations, and 7 Central Valley populations) sampled in the present study and in Berg et al. (2010).

Third, we compared the genetic diversity of the Conaway Ranch Tricolored Blackbird population with that of *Agelaius* congeners, using 3 populations of Red-winged Blackbirds and one population of Yellow-shouldered Blackbirds. This comparison required us to use a subset of the genotypes in the current study, because only 7 loci (LTMR6, Qm10, Dpµ16, Pca3, Ap79, Ap107, and Ap144) had been used to genotype all populations. Of the 2 remaining loci, Ap38 had not been analyzed in Red-winged Blackbirds, and Ap146 was not polymorphic in Yellow-shouldered Blackbirds.

The 3 Red-winged Blackbird populations comprised 2 continental populations and one island population. As continental populations are genetically indistinguishable from each other (Ball et al. 1988; Liu et al. 2015), we used the 2 populations in Liu et al. (2015) with the largest sample sizes (Pennsylvania and Michigan, sampled by I.A.L. in 2005 and S. Lüpold in 2009, respectively). These populations have the highest levels of genetic diversity and effective population sizes of all the reference populations (Pennsylvania, N_e_ ∼430; Michigan, N_e_ ∼infinity, either due to sampling error or a truly large population size not experiencing loss of heterozygosity) (Liu 2015). The third population was a sedentary population of Red-winged Blackbirds on Grand Bahama Island, sampled in 2011, which had lower genetic diversity and a lower effective population size (N_e_ ∼300) than the continental populations (Liu et al. 2015; Liu 2015). The population of Yellow-shouldered Blackbirds, an endangered island endemic, was sampled in Puerto Rico in 2012. This population had the lowest genetic diversity and effective population size (N_e_ ∼70) of all the reference populations (Liu 2015).

We used the bootstrap resampling approach described in Liu et al. (2015) to estimate sample-size-adjusted population allelic diversity, Shannon diversity, and expected heterozygosity across the 5 populations. For each population, we used custom scripts in R 3.1.3 (J. Johndrow personal communication; R Core Team 2015) to take 1000 resamples and calculate the 3 measures for each resample. The populations with the smallest sample sizes were the Conaway Ranch Tricolored Blackbird population (*n* = 50) and the Michigan Red-winged Blackbird population (*n* = 51). Therefore, we used a resample size of 48 individuals so that estimates of uncertainty could be generated for all populations. Values reported are mean ± SD. We used Tukey HSD tests to detect significant pairwise differences across the 5 populations. We then calculated the *P* value as the proportion out of 1000 iterations for which genetic diversity or expected heterozygosity in the Conaway Ranch Tricolored Blackbird population was highest of all populations.

We performed an additional analysis with the cross-species data set using the jackmsatpop function of the R package PopGenKit 1.0 (Paquette 2012) to generate a rarefaction curve of mean raw allelic diversity. This function uses jackknife resampling to measure the number of sampled alleles for a given constant increase in sample size for each population. Although the curves do not give statistical information, they show whether sampling was sufficient to capture population allelic diversity. For each population, we ran 100 repetitions using a stepwise increase of one individual up to that population’s sample size of adults, as in Liu et al. (2015).

## RESULTS

### Microsatellite Quality

Micro-Checker found no evidence for null alleles or dropout in any of the 5 loci tested. Genepop detected linkage disequilibrium between Dpµ16 and Pca3, but the association did not remain significant after a Bonferroni correction. All loci were found not to deviate significantly from Hardy-Weinberg equilibrium (*P* > 0.05).

### Genetic Diversity

The GenAlEx-derived measurements of genetic diversity for the Conaway Ranch Tricolored Blackbird population are shown in Table 1A. Raw allelic diversity of the 9 loci in the Conaway Ranch Tricolored Blackbird population was similar to that of the 8 loci used in the southern California and Central Valley Tricolored Blackbird populations in Berg et al. (2010) (*F*_2,22_ = 0.3, *P* = 0.71, Table 2). Likewise, the 2 sets of loci did not differ significantly in observed or expected heterozygosity (H_o_: *F*_2,22_ = 2.1, *P* = 0.14; H_e_: *F*_2,22_ = 0.6, *P* = 0.57). Thus, genetic diversity was similar across Tricolored Blackbird populations, despite the use of different loci in the present study and Berg et al. (2010).

**Table 1.** GenAlEx measures of sample size (N) and means for raw (N_a_) and effective (N_ea_) number of alleles, Shannon diversity index (I), observed (H_o_) and expected (H_e_) heterozygosity, and inbreeding coefficient (F_IS_) for (**A**) 9 microsatellite loci in Conaway Ranch Tricolored Blackbirds (TRBL_CR), and (**B**) 7 loci in Michigan, Pennsylvania, and Bahamas Red-winged Blackbirds (RWBL); Conaway Ranch Tricolored Blackbirds; and Yellow-shouldered Blackbirds (YSBL). Numbers in parentheses are standard error.

| | N | $N_a$ | $N_{ea}$ | I | $H_o$ | $H_e$ | $F_{IS}$ |
| --- | --- | --- | --- | --- | --- | --- | --- |
| TRBL_CR | 50 | 9.89 | 4.82 | 1.68 | 0.76 | 0.74 | -0.03 |
|  |  | (1.34) | (0.82) | (0.16) | (0.04) | (0.04) | (0.02) |

(B)
| | N | $N_a$ | $N_{ea}$ | I | $H_o$ | $H_e$ | $F_{IS}$ |
| --- | --- | --- | --- | --- | --- | --- | --- |
| RWBL_MI | 51 | 18.71 | 10.21 | 2.46 | 0.83 | 0.88 | 0.057 |
|  |  | (2.79) | (1.83) | (0.18) | (0.03) | (0.03) | (0.025) |
| RWBL_PA | 60 | 19.57 | 10.11 | 2.46 | 0.87 | 0.88 | 0.012 |
|  |  | (3.19) | (1.91) | (0.19) | (0.02) | (0.02) | (0.021) |
| RWBL_Bah | 66 | 9.57 | 5.38 | 1.64 | 0.71 | 0.72 | 0.011 |
|  |  | (2.81) | (1.57) | (0.28) | (0.06) | (0.06) | (0.02) |
| **TRBL_CR** | 50 | 10.29 | 4.71 | 1.68 | 0.74 | 0.73 | -0.013 |
|  |  | (1.41) | (0.97) | (0.18) | (0.05) | (0.05) | (0.026) |
| **YSBL** | 63 | 6.14 | 4.13 | 1.36 | 0.66 | 0.66 | 0.011 |
|  |  | (1.75) | (1.07) | (0.25) | (0.07) | (0.06) | (0.019) |

**Table 2.** Raw allelic diversity (N_a_) and observed (H_o_) and expected (H_e_) heterozygosity of different Tricolored Blackbird populations, using 9 microsatellite loci in the Conaway Ranch population (*n* = 50) and 8 loci in southern California (*n* = 95, 4 populations) and Central Valley (*n* = 122, 7 populations). All data were calculated in GenAlEx. Data for the southern California and Central Valley populations are used with permission and unmodified from Table 3 in Berg et al. (2010). Dpµ16 is the single locus used in both studies.

|  | Conaway Ranch |  |  |  | Southern California |  |  |  | Central Valley |  |  |
| --- | --- | --- | --- | --- | --- | --- | --- | --- | --- | --- | --- |
| Locus | N<sub>a</sub> | H<sub>o</sub> | H<sub>e</sub> | Locus | N<sub>a</sub> | H<sub>o</sub> | H<sub>e</sub> | Locus | N<sub>a</sub> | H<sub>o</sub> | H<sub>e</sub> |
| LTMR6 | 9 | 0.600 | 0.657 | Ase18 | 15 | 0.681 | 0.748 |  | 12 | 0.694 | 0.676 |
| Dpμ16 | 10 | 0.820 | 0.741 | Dpμ16 | 13 | 0.630 | 0.799 |  | 11 | 0.610 | 0.721 |
| Pca3 | 9 | 0.640 | 0.605 | Dpμ1 | 3 | 0.189 | 0.223 |  | 2 | 0.180 | 0.178 |
| Qm10 | 9 | 0.760 | 0.807 | QmAAT10 | 8 | 0.784 | 0.794 |  | 7 | 0.694 | 0.738 |
| Ap38 | 4 | 0.720 | 0.670 | Gf12 | 18 | 0.843 | 0.908 |  | 17 | 0.873 | 0.911 |
| Ap79 | 6 | 0.600 | 0.581 | Lswμ7 | 5 | 0.374 | 0.514 |  | 3 | 0.344 | 0.485 |
| Ap107 | 18 | 0.880 | 0.891 | Gf5 | 10 | 0.533 | 0.630 |  | 5 | 0.504 | 0.573 |
| Ap144 | 11 | 0.900 | 0.856 | Dca32 | 11 | 0.778 | 0.855 |  | 11 | 0.762 | 0.846 |
| Ap146 | 13 | 0.920 | 0.864 |  |  |  |  |  |  |  |  |
| Mean | 9.89 | 0.760 | 0.741 |  | 10.38 | 0.602 | 0.684 |  | 8.50 | 0.583 | 0.641 |

GenAlEx- and bootstrap-derived summaries of genetic diversity for the 5 *Agelaius* populations in the cross-species analysis are shown in Table 1B and Table 3, respectively. The bootstrap found that, out of 1000 resamples, in no iteration did the Conaway Ranch Tricolored Blackbird population have the greatest allelic diversity, Shannon diversity, or expected heterozygosity relative to the other populations (*P* < 0.001, Table 2, Figure 1A). Instead, the continental Red-winged Blackbird populations consistently had the highest values. The profile of the Conaway Ranch Tricolored Blackbird population was most similar to that of the Bahamas Red-winged Blackbird population, although the former had significantly higher values for all measures (*P* for all pairwise comparisons < 0.01, Figure 1B). Lowest for all measures was the Yellow-shouldered Blackbird, which is expected given the low effective population size and endangered status of the species (Liu 2015). These rankings were supported by the rarefaction curve, which predicted that raw allelic diversity in the Conaway Ranch Tricolored Blackbird population approached a plateau above those of the Bahamas Red-winged Blackbird and Yellow-shouldered Blackbird populations but well below the plateau of the continental Red-winged Blackbird populations (Figure 1B).

**Table 3.** Bootstrap-derived measures of mean allelic diversity, Shannon diversity (I), and expected heterozygosity (H_e_) across 7 microsatellite loci in 3 *Agelaius* species, using a resample size of 48 in each of 1000 iterations. Numbers in parentheses are standard deviation. See Table 1 for explanation of abbreviations.

|  | **N** | **Allelic**<br>**diversity** | **I** | **$H_e$** |
| --- | --- | --- | --- | --- |
| **RWBL_MI** | 51 | 16.04<br>(0.56) | 2.36<br>(0.036) | 0.87<br>(0.007) |
| **RWBL_PA** | 60 | 16.25<br>(0.65) | 2.35<br>(0.036) | 0.87<br>(0.006) |
| **RWBL_Bah** | 66 | 8.47<br>(0.31) | 1.59<br>(0.030) | 0.71<br>(0.012) |
| **TRBL_CR** | 50 | 9.03<br>(0.4) | 1.63<br>(0.036) | 0.73<br>(0.011) |
| **YSBL** | 63 | 5.79<br>(0.18) | 1.33<br>(0.023) | 0.66<br>(0.011) |

**Figure 1.**
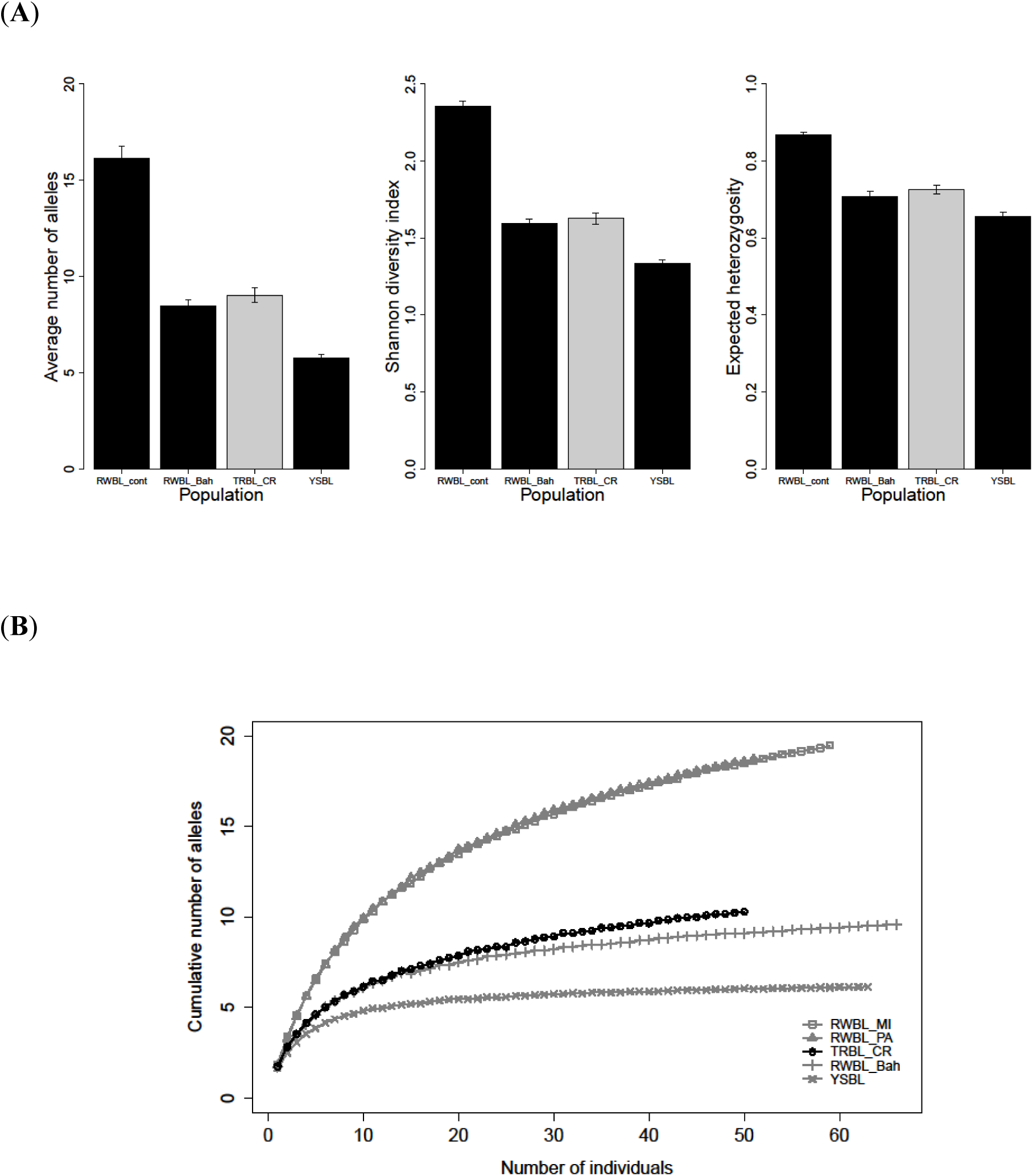
(**A**) Genetic diversity estimated from bootstrap simulations using 7 microsatellite loci across 3 species. “RWBL_cont” refers to the mean measures of 2 continental (Michigan and Pennsylvania) Red-winged Blackbird populations. “RWBL_Bah” refers to a Bahamas Red-winged Blackbird population. “YSBL” refers to a Yellow-shouldered Blackbird population. The Conaway Ranch Tricolored Blackbird population (TRBL_CR) has significantly less genetic diversity than the continental populations of Red-winged Blackbirds and similar diversity to the Bahamas population of Red-winged Blackbirds. (**B**) Rarefaction curve, using mean raw allelic diversity of 7 microsatellite loci, showing the genetic diversity of the Conaway Ranch Tricolored Blackbird population (black circles) relative to other *Agelaius* species. The Conaway Ranch Tricolored Blackbird population approaches a maximum similar to that of the Bahamas Red-winged Blackbirds population (gray vertical lines).

### Inbreeding coefficient

The mean inbreeding coefficient (F_IS_) for the Conaway Ranch Tricolored Blackbird population was negative (F_IS_ = −0.03, Table 3). This result contrasted with the moderately high inbreeding coefficients for the southern California and Central Valley Tricolored Blackbird populations reported in Berg et al. (2010) (F_IS_ = 0.121 and 0.090, respectively).

## DISCUSSION

The Tricolored Blackbird has experienced severe declines in total population size over the last century due to destruction of native habitats and large-scale, chronic nesting failures (Beedy 2008; Meese 2013; Meese et al. 2014). The species is currently under review to be listed as Endangered under both the California and U.S. Endangered Species Acts. Our analysis revealed that the genetic diversity of a Central Valley population is comparable to that of a small, isolated island population of the related Red-winged Blackbird. Because of this finding, the results in our study and in Berg et al. (2010) collectively support both state and federal protection of the Tricolored Blackbird to maintain existing gene flow and genetic diversity.

### Low Genetic Diversity

We recognize that a single-population study yields limited insight into the conservation measures required for a whole species. Such studies nevertheless provide valuable data by complementing existing demographic literature and enabling policymakers to better assess the degree and immediacy of the threats facing a species. To maximize the useful inferences that can be drawn from our findings, we frame our results relative to estimates reported in other Tricolored Blackbird populations and in populations of *Agelaius* congeners. We also discuss our results in the context of known extinction risk factors and recommend future work, stemming from our study, whose findings will be informative for managers.

We found that the genetic diversity of the Conaway Ranch Tricolored Blackbird population is more similar to that of a small, non-migratory Red-winged Blackbird population in the Bahamas than to Red-winged Blackbird populations in the continental U.S. By extension, this relatively low diversity appears to be shared across all studied populations, given that the genetic profile of the Conaway Ranch population was similar to that of the 11 populations in Berg et al. (2010) (Table 2). It is possible that the Tricolored Blackbird’s genetic diversity always has been comparably low, because its narrow geographic range might have imposed upper limits on its total population size and thus its genetic diversity (e.g., Frankham 1997; Hague and Routman 2016). Genotypes from historical specimens collected over different time points (which Berg et al. (2010) report are available for this species dating back to 1861) would be helpful to determine whether genetic diversity previously approached that of continental Red-winged Blackbird populations. Paired with census data, historical specimens also would be useful in testing the relationship between genetic diversity and population size and connectivity (e.g., Pacioni et al. 2015). At present, the concern is that a population of a species historically known for its abundance (Neff 1937) currently displays levels of genetic diversity resembling that of an isolated *Agelaius* blackbird population.

Genetic effects of range limits potentially can be buffered by connectivity between colonies (e.g., Castellanos-Morales et al. 2015). Indeed, a promising finding is that Tricolored Blackbird populations currently do not seem to be isolated, as no structure was found within or across southwest California and Central Valley populations (Berg et al. 2010). Precise rates of migration between specific populations are unknown, but a limited number of recaptures and sightings of banded birds reveal that small numbers of birds move between southern California and the Central Valley (R.J. Meese personal observation). It is not clear whether this movement is facilitated by habitat connectivity between colonies, or whether individuals are simply able to fly over unsuitable habitat en route to a different colony. However, evidence of dwindling site occupancy and recruitment due to habitat loss (Holyoak et al. 2014) suggests migration may become more difficult with increasing or prohibitive distances between colonies. Thus, continuing habitat loss could lead (directly or indirectly) to population fragmentation across the Tricolored Blackbird’s range, accelerating declines in genetic diversity due to absolute colony loss. Given the already high rates of population size reduction and the risk of further decreases in genetic diversity, existing gene flow across colonies will be critical to monitor and preserve.

In comparing sequence-level differences between 10 Tricolored and 31 continental Red-winged blackbirds, Barker et al. (2012) found Tricolored Blackbirds had less polymorphic mtDNA, according to summary statistics such as nucleotide diversity (π) and Watterson’s estimator (θ). Although different methods (summary statistics vs. likelihood and Bayesian analyses) led to different inferences of the demographic history of these species, DNA-sequence polymorphisms were consistently lower in Tricolored Blackbirds. These results align with the microsatellite data in showing that contemporary Tricolored Blackbirds are less genetically diverse than continental populations of the closely related Red-winged Blackbird.

### Low Inbreeding Coefficient

In contrast to the findings of Berg et al. (2010) across 11 Tricolored Blackbird populations, we did not find evidence of inbreeding in the Conaway Ranch population. These disparate results could be due to temporal variation in inbreeding caused by differences in demography or mate choice. For example, inbreeding coefficients could be higher in the other populations because those populations contained more related individuals or dispersal across them was more limited. As a result, Conaway Ranch as a single population may have had lower measures of inbreeding than the mean values found for the southern California and Central Valley populations. Alternatively, because signatures of inbreeding (namely loss of heterozygosity) require multiple generations to manifest, inbreeding in the Conaway Ranch population may not have been immediately apparent if losses in this colony were recent.

Our findings suggest that individuals in the Conaway Ranch population are, on average, not breeding with close relatives. However, the recorded decline in census population size, combined with the relatively low levels of genetic diversity, signal that inbreeding and its accompanying fitness consequences remain risks both to the Conaway Ranch and the other surveyed populations. In addition to the yearly censuses already in place, we recommend that further measures of inbreeding coefficients be taken to reconcile the calculations in the 2 studies, and that longitudinal surveys of specific fitness components (e.g., offspring viability, first-year survival, and reproductive success) be conducted in selected populations to evaluate the presence of inbreeding depression. Moreover, the larger sample sizes obtained from these measurements could give informative estimates of effective population size—which our smaller data set was unable to provide—and thus predict the risk of mutation and genetic load (Lynch et al. 1995). Support granted by legal protections would help enable these more intensive surveys and provide the opportunity to test whether survival and/or fecundity have been affected by habitat loss and population size decline.

### Conservation Implications

The results in our study and in Berg et al. (2010) support the need for legal protections of the Tricolored Blackbird. Current losses of breeding and foraging habitat, long-term destruction of some of the largest colonies in grain fields, incidental shooting, and elevated rates of predation and pesticide exposure have led to rapid and ongoing reductions in total population size. The blackbird’s restricted range and decreased genetic diversity may accelerate the effects of drift, preventing immediate fitness recovery while also weakening the species’ ability to respond over time to environmental change (Lande 1988; Reed and Frankham 2003).

Additionally, the blackbird’s colonial nature increases its vulnerability both to large-scale losses and to density-dependent social effects (i.e. Allee effects). Colony size is a predictor of adult survival in Tricolored Blackbirds (Weintraub and George 2012) and, in other birds, correlates positively with dispersal and negatively with predation (Serrano et al. 2005; but see Brown et al. 2016). The 75% decrease in the size of the largest colony over the last decades suggests colonies could be at risk for demographic instability if they are not adapted to living in present-day group sizes. Together, all of the above factors expose the Tricolored Blackbird to greater risk of extinction than related species (such as the Red-winged Blackbird) with large ranges and population sizes, non-colonial life histories, and multiple sources of genetic variation.

The well-documented population decline and the comparatively low genetic diversity reported here indicate that protections provided by a listing under the ESA and CESA are warranted. These further protections would help ensure effective conservation actions by providing management plans, legal enforcement, and federal aid to fill gaps in scientific knowledge, some of which have been outlined here. The immediacy and severity of the threats facing the Tricolored Blackbird, coupled with the genetic data presented here, justify further candidate assessment and additional legal protections.

## ACKNOWLEDGMENTS

We thank Steve Nowicki and Mohamed Noor at Duke University for their guidance and the use of lab space, and Jill Soha for her comments on the manuscript. We are grateful to Mike Hall for his generous support of scientific research at the Conaway Ranch. Field protocols were approved by Duke University’s Institutional Animal Care and Use Committee and all relevant local and national institutions. Fieldwork was conducted in compliance with the *Guidelines to the Use of Wild Birds in Research*. This work was funded by a National Science Foundation Dissertation Improvement Grant (award no. IOS-1110782), a Sigma Delta Epsilon Fellowship from Graduate Women in Science, and the Office of the Provost at Duke University to I.A.L. Funders had no input into the content of the manuscript, nor was funder approval required of the manuscript before submission or publication. I.A.L. conceived the idea, collected the data, performed the analyses, and wrote the paper. R.J.M. contributed expertise in Tricolored Blackbird biology, provided guidance in the field, and edited the paper.

