## Supplementary material for "Low genetic diversity in a population of Tricolored Blackbird (*Agelaius tricolor*), a species pending Endangered status": Table S1

Table S1. Primers used to genotype Tricolored Blackbirds. *L* = allele length, *n* = number of alleles.

| Locus | Forward/reverse primer sequences | Repeat motif | *L* | *n* | GenBank | Reference |
|---|---|---|---|---|---|---|
| LTMR6 | GCCATGCCACAGGAGTGAGTC | GT | 183-202 | 10 | FM201465.1 | McDonald and Potts (1994) |
| | AGTCATCTCCATCMGGGCAT | | | | | |
| Qm10 | GGAATTCCAGTATGTGAATGAGTC | AAT | 210-238 | 9 | AF013235.1 | Hughes et al. (1998) |
| | ATTGCAAAAAACAGAAGCATTTTAAC | | | | | |
| Dpμ16 | ACAGCAAGGTCAGAATTAAA | AC,GC | 146-165 | 11 | AM262982.1 | Dawson et al. (1997) |
| | AACTGTTGTGTCTGAGCCT | | | | | |
| Pca3 | GGTGTTTGTGAGCCGGGG | GT | 153-178 | 9 | AJ279805.1 | Dawson et al. (2000) |
| | TGTTACAACCAAAGCGGTCATTTG | | | | | |
| Ap38 | GGAGGGAGACCTCTTAATAC | CATC | 237-257 | 6 | JF907496.1 | Barker et al. (2011) |
| | CGACAGAGCTGGTGTCAAAA | | | | | |
| Ap79 | CCACTTCTGCTGAACATAGGG | AAC | 218-241 | 6 | JF907499.1 | Barker et al. (2011) |
| | GTGCTGCAATTGTGGTCTTG | | | | | |
| Ap107 | GAAACATCCAAACCTGGCTTG | AGAT | 211-255 | 19 | JF907500.1 | Barker et al. (2011) |
| | AATGGACGTGCAGCCCTTC | | | | | |
| Ap144 | TCCATAACACAGTTGTCAGAG | AGAT | 200-241 | 12 | JF907502.1 | Barker et al. (2011) |
| | CTTACACAGGCACACAAACC | | | | | |
| Ap146 | ACATTCCCAGGTCTCACTGC | AAC | 112-152 | 14 | JF907503 | Barker et al. (2011) |
| | GTTCACGGATCGGAAAGAATC | | | | | |
